# Cost-effective genomic prediction for fertility traits: A comparison of machine learning and conventional models using low-coverage sequencing in Holstein heifers

**DOI:** 10.1101/2025.08.01.668144

**Authors:** Z.B. Zhang, A. Wang, Q.Y. Wang, S.Q. Gao, L.L. Wang, H.H. Hu, H.A Nanaei, A.M. Shah, G.L. Liu, K. Zhu, X.Z. Lv, R. Li, Y. Jiang

## Abstract

Fertility is one of the major factors affecting the efficiency of dairy herd, and genomic selection (**GS**) on milk yield, while ignoring fertility, has resulted in a decline in heifer fertility. Consider the cost of breeding, it is important to enhance the accuracy of GS with cost-effective way for fertility traits. This study investigates the genomic prediction (**GP**) of fertility traits in Holstein heifers using both machine learning (**ML**) and conventional methods, based on data from SNP arrays and low-coverage sequencing data. In this study, we collected 45,320 Holstein heifers with phenotype and pedigree records, from which we generated genomic data for 3,683 Holstein heifers using lcGWS. We first estimated the heritability for age at first service (**AFS**), gestation length (**GLh**) and age at first calving (**AFC**). We then compared the prediction performance of ML methods, kernel ridge regression (**KRR**), support vector regression, and random forest regression, with GBLUP, ssGBLUP and BayesR3 regarding GP accuracy and unbiasedness. Inputs for ML includes genomic relationship matrices (**GRM**), principal components, and SNPs. The results revealed that the heritability for the three fertility traits ranged from 0.09 to 0.48. Prediction accuracy from imputed low-coverage sequencing data was comparable to that from standard SNP chips. When both pedigree and genotypic data were used for GS, ssGBLUP yielded the highest prediction accuracy for AFS and GLh. Crucially, using only genomic data, KRR_GRM improved GP accuracy by up to 28.57% compared to GBLUP and by up to 9.46% compared to BayesR3. Our results highlight the effectiveness of low-coverage sequencing data in breeding applications and the ML’s potential to enhance GP accuracy for fertility traits, offering practical insights for dairy breeding programs.

**Implications:** To improve the accuracy of GS with cost-effective way for fertility traits, this study confirms that low-coverage sequencing offers both accuracy and cost-effectiveness for fertility in Holstein heifers. The research provides a decision-making framework for breeding workers: the ssGBLUP model is optimal when combining pedigree data, while machine learning methods are superior with only genomic data. This study offers a practical tool for achieving efficient and economical genetic improvement in dairy cattle.

## Introduction

Many economic traits are controlled by genes with small effects (Zhang et al., 2019), so a large amount of data is required to estimate the effect of SNPs accurately. The genomic selection (**GS**), as proposed by Meuwissen et al (Meuwissen et al., 2001), assumes that the genetic variants are in linkage disequilibrium (**LD**) with the quantitative trait loci (**QTL**). The models used for GS are usually classified into best linear unbiased prediction (**BLUP**) and Bayesian models. Normally, GBLUP (VanRaden et al., 2009) is faster and relatively accurate. The Bayesian methods such as BayesA (Meuwissen et al., 2001), BayesB (Meuwissen et al., 2001), BayesCπ (Habier et al., 2011), BayesDπ (Habier et al., 2011) and BayesR (Erbe et al., 2012) are highly accurate but require estimating all SNP effects simultaneously, resulting in lower computational efficiency. A type of block Gibbs sampling for estimating Markov chain SNP effects has been introduced recently by BayesR3. It dramatically reduces the running time by iteratively sampling each SNP effect n times from the conditional block posterior (Breen et al., 2022).

Considering the cost of breeding, obtaining genome-wide variation at a low cost is essential for dairy cattle breeding. The advent of low-coverage sequencing offers a viable solution to these challenges. The low-coverage sequencing has been successfully used to find key genes in humans (Martin et al., 2021) or decrease the cost of breeding in pigs (Yang et al., 2021), chicken (Ronneburg et al., 2023) and aquatic animals (Yang et al., 2024). Our previous work (Zhang et al., 2023) established the first publicly available reference panel for cattle and optimized the imputation workflow for low-coverage sequencing, the imputation accuracy for Holstein was more than 99%. Therefore, low-coverage sequencing is a high-throughput and cost-effective genotyping method that warrants further application in dairy cattle.

The first species to adopt GS was dairy cattle, and studies have revealed that GS breeding cost of dairy cows was 92% less than traditional progeny testing (Schaeffer, 2006). In the United States, the genetic gain for annual yield traits in Holstein cows has increased from approximately 50% to 100% (Garcia-Ruiz et al., 2016). Fertility is one of the major factors affecting the efficiency of dairy herd, and GS on milk yield with ignoring fertility has resulted in a decline in heifer fertility (Alves et al., 2023; Aponte et al., 2024). Importantly, since the heritability of some fertility traits is relatively low, improving the accuracy of GS for heifer fertility traits is of importance.

The machine learning (**ML**) methods are flexible and powerful, capable of uncovering hidden data patterns in large datasets with hundreds or thousands of potential explanatory variables (Chafai et al., 2023). These methods play an important role in improving GS accuracy (Ehret et al., 2015; An et al., 2021; Liu et al., 2022; Chafai et al., 2023; Lee et al., 2023; Liang et al., 2023; Xiang et al., 2023; Yan and Wang, 2023). A previous study also demonstrated the ML method improved the accuracy of fertility traits by 15-20% compared to GBLUP and Bayesian methods in 2,566 Chinese Yorkshire pigs (Wang et al., 2022). Compared to GBLUP, ML methods have shown a 33% improvement in GP accuracy for lifetime productivity and a 16% improvement for lifetime litter size in pigs (Hong et al., 2024). Thus, ML may play an important role in improving GP accuracy for fertility traits in livestock.

This study first estimated the variance components of fertility traits using pedigree data in Holstein heifers. We also compared the accuracy of different SNP arrays on GP accuracy with low-coverage sequencing data and the prediction performance of various ML methods on fertility traits of heifers. These findings demonstrated the effectiveness of lcGWS data in breeding applications and provide valuable insights for enhancing GP accuracy in heifers.

## Material and methods

### Animal and genotype data

In this study, we collected fertility traits and pedigree of 45,320 Holstein heifers, and whole blood samples from 3,683 heifers were also collected during the routine health checks from farms. DNA was extracted and sent to MolBreeding Biotech Co. Ltd (Shijiazhuang, Hebei) for low-coverage sequencing. The obtained raw reads were subjected to quality control using fastp v0.20.0 (Chen et al., 2018) with default parameters. Subsequently, the reads were aligned to the bovine reference genome ARS-UCD1.2 using BWA MEM v0.7.17 (Li and Durbin, 2010). The bam file conversion, sorting, and duplicate read filtering were performed using samtools v1.10 (Li et al., 2009) and Picard v2.20.1 (http://broadinstitute.github.io/picard/). The average sequencing depth was 1.

Referring to the reference panel and imputation workflow from the previous study (Zhang et al., 2023), the obtained low-coverage sequencing data was imputed to the whole-genome level. As a preprocessing step for genotype data, firstly, we filtered the imputed SNP set with an LD threshold of r^2^ > 0.9 and a minor allele frequency (MAF) < 0.01, resulting in 5,559,783 SNPs, referred to as 5M in this study. Additionally, we extracted 46,089, 127,582, and 616,134 SNPs from the imputed SNP set based on the BovineSNP50 v3 BeadChip, GGP BeadChip GGPHDV3 and SNP BovineHD BeadChip, respectively, referred to as 50K, 140K, and 777K.

Following the approach from previous studies (Liang et al., 2021; Wang et al., 2023a), we used GCTA v1.94.1 (Yang et al., 2011) to calculate the genomic relationship matrix (**GRM**), and PLINK v1.90b7 (--pca) (Chang et al., 2015) to compute and extract the principal components (**PCs**) which explained top 95% of feature variance. In this study, we generated three forms of data, SNP, PCs and GRM, based on genotype information to serve as input features for machine learning methods.

### Phenotype and fixed effect correction

In our study, we utilized three fertility-related traits from Holstein heifers: age at first service (**AFS**), gestation length in days of heifer (**GLh**) and age at first calving of heifer (**AFC**). Additionally, to ensure comparability among different models in GP, we remove the fixed effects in prediction for each trait, the fixed effects used are shown in Supplementary Table S1.

### Statistical models

The linear mixed model was used for variance components estimation. Three conventional methods (GBLUP、ssGBLUP and BayesR3) and three ML methods, including kernel ridge regression (**KRR**), support vector regression (**SVR**) and random forest regression (**RFR**), were used to perform GS of fertility traits. We defined the corrected phenotype as y_c_ and the predicted phenotype (or GEBV for conventional method) as *y*_p_.

### Variance components estimation

The model used for variance component estimation is as follows:

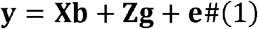

In this equation, **y** is the vector of phenotypes, **X** is the incidence matrix; **b** is the vector of fixed effects as showed in Supplementary Table S1, **z** is the incidence matrix, **g** is the vector of random additive genetic effects, **e** is the vector of random residual effects. The random effects are distributed as follows: 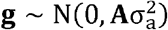 and 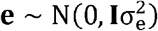, where **A** is the additive relationship matrix, and 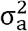 and 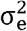 represent the additive genetic variance and residual variance, respectively. The variance component estimation was performed in BLUPF90(Misztal et al., 2018).

### GBLUP

The model used for GBLUP is as follows:

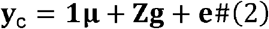

In this equation, **y**_**c**_ is the vector of corrected phenotypes, 1 is a vector of ones, **μ** is the overall mean value, **g** is the vector of random additive genetic effects, **z** is the incidence matrix, and **e** is the vector of random residual effects. The random effects are distributed as follows: 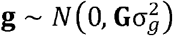 and 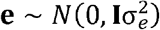, where **G** is GRM, and 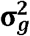 and 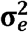 represent the additive genetic variance and residual variance, respectively.

### ssGBLUP

The model used for ssGBLUP is as follows:

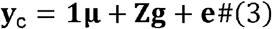

In this equation, **y**_**c**_ is the vector of corrected phenotypes, 1 is a vector of ones, **μ** is the overall mean value, **g** is the vector of random additive genetic effects, **z** is the incidence matrix, and **e** is the vector of random residual effects. The random effects are distributed as follows: 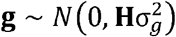 and 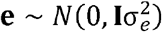, where **H** is an additive kinship matrix constructed by combining pedigree and genotype information, and 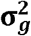 and 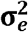 represent the additive genetic variance and residual variance, respectively.

### BayesR3

The formula for Bayesian methods is as follows:

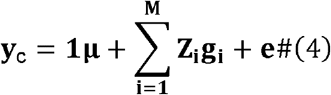

where **y**_**c**_ represents the vector of observations (i.e., corrected phenotypes), 1 is a vector of ones, **μ** is the overall mean value, **z**_**i**_ denotes the genotype at the i-th SNP locus, g_i_ represents the effect value at the i-th SNP locus, **e** denotes the random residual effects, following a distribution 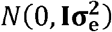. The core distinction among a series of Bayesian methods lies in the rational assumptions made regarding the distribution of marker effects g_i_ and their variance 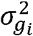.

BayesR (Erbe et al., 2012) categorizes markers into four groups based on the magnitude of their effects: large, medium, small, and null effects. Markers within the same group have the same variance, and there is a consistent gradient ratio between different groups. The specific distribution is as follows:

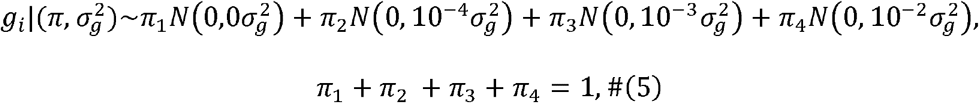

BayesR3 (Breen et al., 2022) is an enhanced version of BayesR that employs a block Gibbs sampling scheme to estimate the effects of Markov chain SNPs. By iteratively sampling each SNP effect n times from the conditional block posterior, BayesR3 significantly reduces running time. In this study, we employed the BayesR3 in R language.

### Kernel ridge regression

KRR (Vovk, 2013) is used to establish a nonlinear relationship model between input features and corresponding target variables. It extends ridge regression by incorporating kernel functions to handle nonlinear relationships. The fundamental idea behind KRR is to introduce the kernel trick on top of ridge regression. It begins by mapping the input features through a nonlinear mapping function (kernel function) into a high-dimensional feature space and performing linear regression in that feature space. By utilizing the kernel function without explicitly computing the high-dimensional feature space, KRR can create a nonlinear model in a low-dimensional input space. The prediction function for KRR can be expressed as:

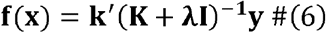

Here, **K** is the kernel matrix, and **I** is the identity matrix, λ is the regularization constant. The most suitable kernel function and regularization strength were determined by grid search.

### Support vector regression

SVR (Drucker H, 1997) is an extension of support vector machines applied to regression problems. The basic principle of SVR is to map the input data from the original space to a feature space using a nonlinear kernel function, where modeling and prediction occur. The training objective of SVR is to establish a hyperplane in the feature space that minimizes the distance to the farthest data points from the hyperplane while also minimizing the within-class variance of all sample data. In other words, SVR aims to find a hyperplane that maximizes the margin while allowing a certain degree of error for each training sample. The hyperplane is defined by a subset of the training samples, called support vectors, which are the closest data points to the hyperplane. The prediction is made by evaluating the distance of the test sample to the hyperplane in the feature space. The SVR problem was formalized as:

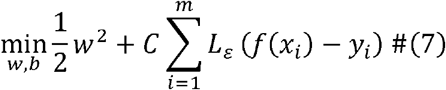

Here, C is the regularization constant,*L*_*ε*_ is the *ε*-insensitive loss:

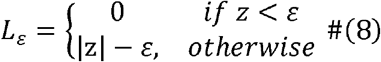

The final prediction model was as followed:

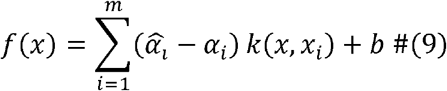

Here, *k*(*x*_*i*_, *x*_*j*_) = *ϕ* (*x*_*i*_)^*T*^ *ϕ* (*x*_*j*_) is the kernel function. The most suitable kernel function was determined by grid search.

### Random forest regression

RFR (Breiman, 2001) is designed based on the ensemble learning technique, known as bagging, which involves combining multiple independent decision trees. Each decision tree in the random forest makes predictions by using a subset of the training samples and features. During both training and testing, each tree can be processed in parallel, enhancing computational efficiency. By aggregating the outputs of multiple decision trees, we can determine the predicted value for the new sample. Combining the predictions from several trees reduces overfitting and increases the overall accuracy and robustness. The predicted value was as follows:

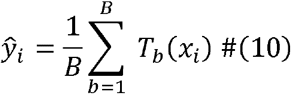

Here, *B* is the number of decision trees in the forest, *T*_*b*_(*x*_*i*_) is an individual regression tree. The depth of the tree was determined by grid search.

The aforementioned ML methods were implemented and tested using the scikit-learn v1.2.2 (Pedregosa et al., 2011) package in Python. In addition, the hyperparameter selection was performed only in the training population, and we tested many different hyperparameters combinations to select the best one. Specific hyperparameters were chosen based on accuracy and runtime using grid search for each method and genotype input format to improve the performance of the ML methods. The detailed parameters can be found in Table 1.

**Table 1:** The selected hyperparameters for ML methods.

| Genotype input format | ML method | Selected hyperparameters |
| --- | --- | --- |
| SNP | KRR | kernel=cosine, alpha=1.15 |
|  | SVR | kernel=poly |
|  | RFR | n_estimators=50 |
| GRM | KRR | kernel=cosine, alpha=6.0 |
|  | SVR | kernel=rbf |
|  | RFR | n_estimators=50 |
| PCs | KRR | kernel=cosine, alpha=6.0 |
|  | SVR | kernel=rbf |
|  | RFR | n_estimators=50 |

Abbreviations: SNP = Single nucleotide polymorphism; GRM = Genomic relationship matrix; PCs = Principal components; KRR = kernel ridge regression; SVR = support vector regression; RFR = random forest regression.

### Metrics to assess GP performance

We evaluated the predictive performance of all methods based on 10-fold cross-validation with 10 repetitions. To ensure the reproducibility of cross-validation, random seed values for the 10 repetitions remained unchanged. We used three metrics based on the validation set within each fold of cross-validation to assess the GP method’s performance.

First, we calculated the Pearson correlation coefficient between y_c_ and y_p_, which have used to evaluated the accuracy of the GP (Yin et al., 2020; Liang et al., 2023; Wang et al., 2024).

Second, we assessed the consistency between the corrected phenotype and the predicted values using prediction unbiasedness (Wang et al., 2022; Wang et al., 2023b). A value closer to 1 indicates a higher consistency between y_c_ and y_p_.

Finally, we measured the running time of each test of ML in seconds to evaluate the computational efficiency of each method under the same hardware environment. All tests were conducted on an 8-core Intel Xeon Platinum 8358 CPU @ 2.60GHz platform.

## Results

### Descriptive statistics

Descriptive statistics for three fertility traits in Holstein heifers are shown in Table 2. We collected phenotypes and pedigrees for AFS, GLh and AFC from 45,320, 36,866, and 36,886 heifers, respectively. The mean values for AFS, GLh and AFC were 445.09, 274.22, and 745.41, respectively. Among these, 3,683 heifers were genotyped.

**Table 2:**
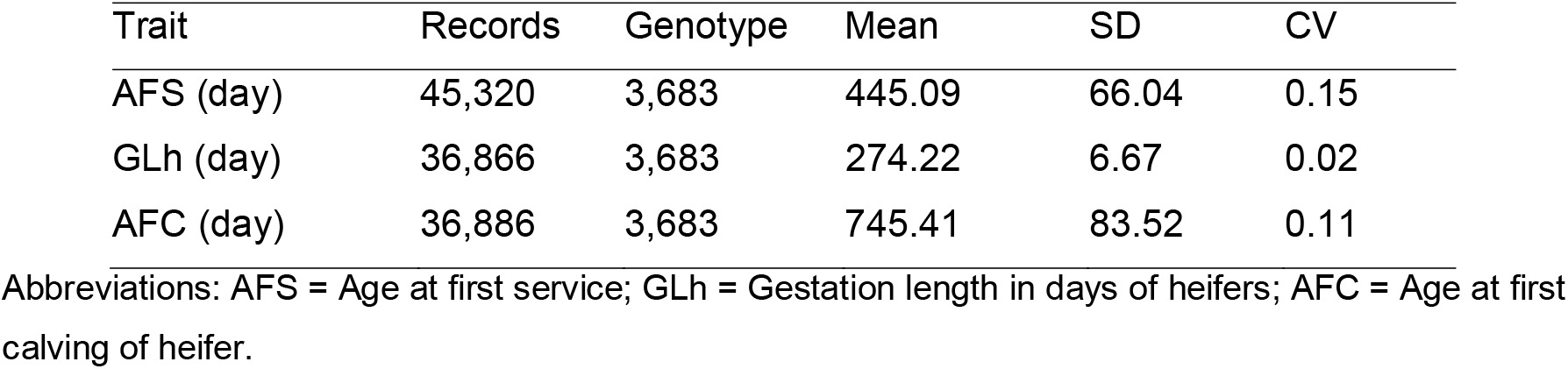
Descriptive statistics of three fertility traits in Holstein heifer.

| Trait | Records | Genotype | Mean | SD | CV |
| --- | --- | --- | --- | --- | --- |
| AFS (day) | 45,320 | 3,683 | 445.09 | 66.04 | 0.15 |
| GLh (day) | 36,866 | 3,683 | 274.22 | 6.67 | 0.02 |
| AFC (day) | 36,886 | 3,683 | 745.41 | 83.52 | 0.11 |

Abbreviations: AFS = Age at first service; GLh = Gestation length in days of heifers; AFC = Age at first calving of heifer.

### The heritabilities of three fertility traits in Holstein heifers

Based on pedigree data, the heritability estimates (***h***^***2***^) calculated for the three fertility traits ranged from 0.09 to 0.47 (Table 3). AFS showed the highest heritability (0.47), AFC had the moderate heritability (0.18), and GLh had low heritability (0.09).

**Table 3:** Estimates of additive genetic variance 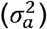, residual variance 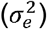, and heritability (*h*^*2*^) for three fertility traits in Holstein heifers.

| Trait | Number | $\sigma_a^2$ | $\sigma_e^2$ | $h^2$ | SE |
| --- | --- | --- | --- | --- | --- |
| AFS (day) | 45,320 | 178.69 | 200.23 | 0.47 | 0.003 |
| GLh (day) | 36,866 | 9.41 | 90.91 | 0.09 | 0.002 |
| AFC (day) | 36,886 | 350.51 | 1632.4 | 0.18 | 0.002 |

Abbreviations: AFS = Age at first service; GLh = Gestation length in days of heifers; AFC = Age at first calving of heifer.

### The impact of SNP density on prediction performance

The prediction accuracy of the three ML input features varied across different SNP densities (Fig. 1). When genotype data was inputted into the ML algorithms as an SNP matrix, KRR and SVR demonstrated an average gain in prediction accuracy of 1.81% and 1.83%, in prediction accuracy at the 140K SNP chip compared to the 50K SNP chip, respectively, while RFR showed an average decrease of 2.26%. At the 777K SNP chip, KRR and RFR exhibited an average increase of 0.23% and 1.05% in prediction accuracy compared to the 140K SNP chip, while SVR showed an average decrease of 0.70%, respectively.

**Fig. 1.**
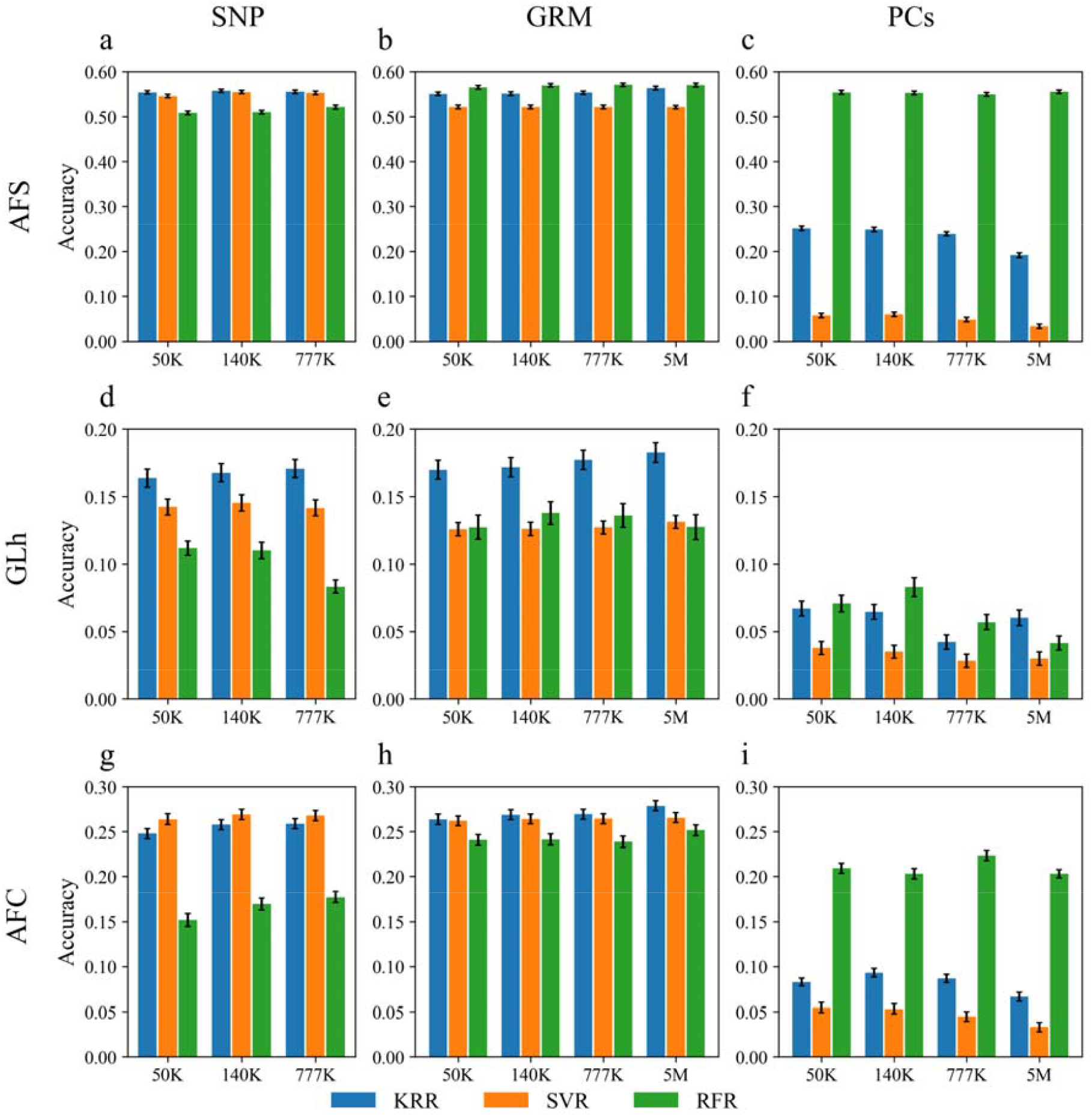
Prediction accuracy of KRR, SVR and RFR. Each row in the figure corresponds to one of the traits: AFS, GLh and AFS. Each column corresponds to one of the inputs: SNP, GRM, and PCs. Each subplot represents the prediction accuracy of three machine learning methods (KRR, SVR, and RFR) at multiple SNP marker densities. (a-c). The prediction accuracy of ML methods for the AFS trait using SNP, GRM, and PCs inputs, respectively. (d-f). The prediction accuracy of ML methods for the GLh trait using SNP, GRM, and PCs inputs, respectively. (g-i). The prediction accuracy of ML methods for the AFC trait using SNP, GRM, and PCs inputs, respectively. The colors blue, orange, and green represent KRR, SVR, and RFR, respectively; the standard error is shown via the error bar, the same as below.

When genotype data was inputted into the ML algorithms as a GRM, KRR, RFR and SVR demonstrated an average gain in prediction accuracy of 0.73%,1.67% and 0.27%, in prediction accuracy at the 140K SNP chip compared to the 50K SNP chip, respectively. At the 777K SNP chip, the KRR and SVR methods showed an average increase of 0.86% and 0.16%, respectively, in prediction accuracy compared to the 140K SNP chip, while RFR showed an average decrease of 0.35%. At the 5M SNP density, KRR, RFR and SVR demonstrated an average gain in prediction accuracy of 2.51%,0.36% and 0.52% respectively, compared to the 777K SNP chip.

When genotype data was inputted into the ML algorithms as PCs, at the140K SNP chip, the prediction accuracy of the KRR and RFR increases by an average of 1.26% and 0.63 respectively compared to the 50K SNP chip, while the SVR methods show an average decrease of 1.36% respectively. At the 777K SNP chip, the prediction accuracy of the KRR, RFR and SVR methods shows an average decrease of 9.49%,1.15% and 18.19% respectively. At the SNP density of 5M, the prediction accuracy of the KRR, RFR and SVR methods shows an average decrease of 13.47%, 3.58% and 20.41% respectively.

Additionally, the efficiency of ML methods was also influenced by the SNP density to some extent, depending on the format of the input data (Fig. 2). Specially, the running time with SNP input increased proportionally with SNP density, whereas the running time with GRM and PCs input remained stable cross a range of marker densities.

**Fig. 2.**
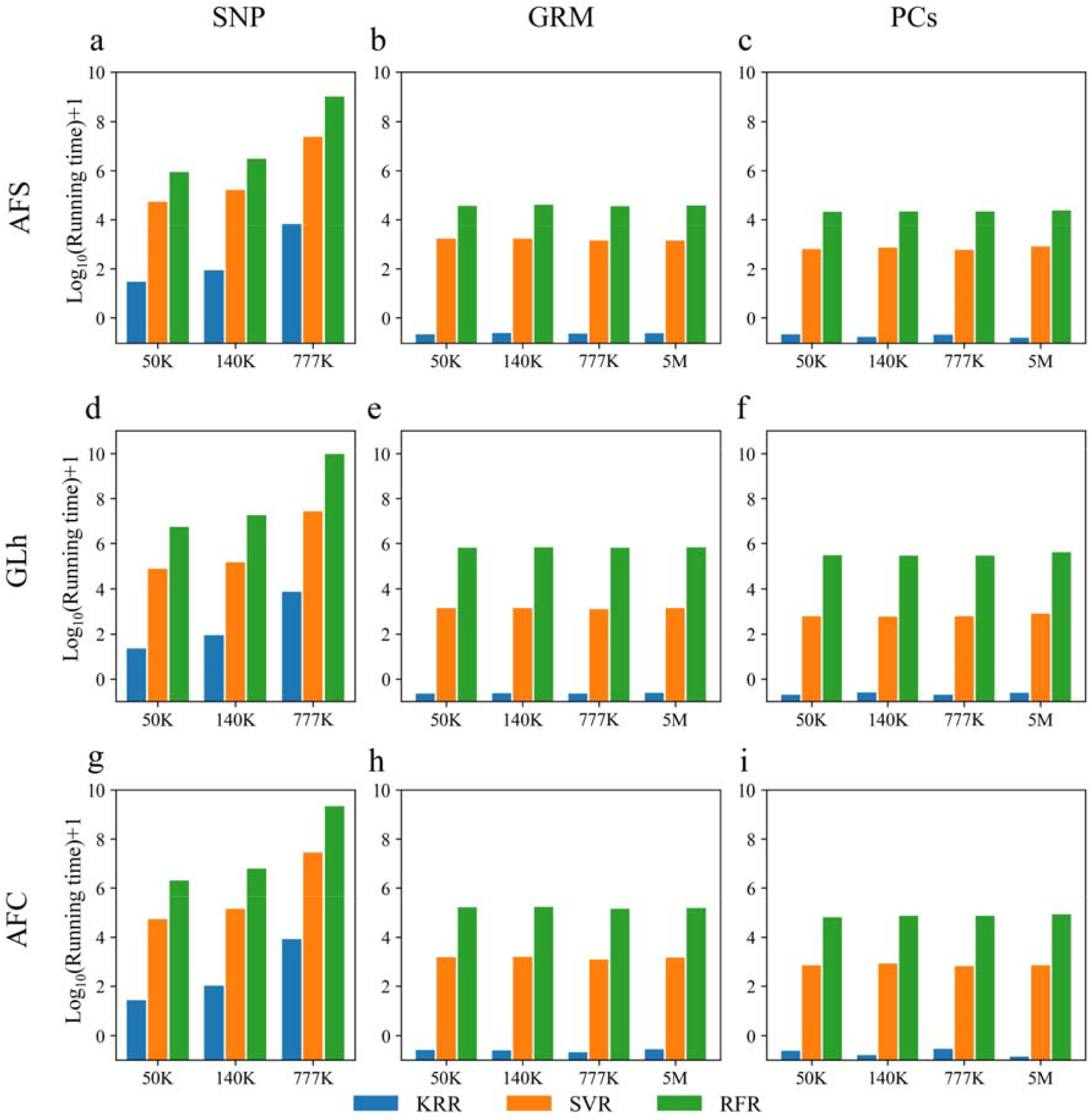
The running time of KRR, SVR and RFR. Each row in the figure corresponds to one of the traits: AFS, GLh and AFS. Each column corresponds to one of the inputs: SNP, GRM, or PCs. Each subplot represents the running time of three machine learning methods (KRR, SVR, and RFR) at multiple SNP marker densities. (a-c). The running times of ML methods for the AFS trait using SNP, GRM, and PCs inputs, respectively. (d-f). The running times of ML methods for the GLh trait using SNP, GRM, and PCs inputs, respectively. (g-i). The running times of ML methods for the AFC trait using SNP, GRM, and PCs inputs, respectively. Notes: The time for GRM and PCs was not included.

For the conventional methods GBLUP, ssGBLUP and BayesR3, the prediction accuracy increased slightly with the increase in SNP density slightly (Table 4). When comparing the prediction accuracy at the 140K SNP density to that at the 50K SNP chip, GBLUP demonstrated an average gain of 2.38%. At the 777K SNP chip, it exhibited an average decrease of 1.16% in prediction accuracy compared to the 140K SNP chip. At the 5M SNP density, GBLUP improved 1.18% in prediction accuracy compared to the 777K SNP chip. About ssGBLUP, when comparing the prediction accuracy at the 140K SNP chip to that at the 50K SNP chip, ssGBLUP demonstrated an average gain of 0.98%. At the 777K SNP chip, it exhibited an average increase of 1.94% in prediction accuracy compared to the 140K SNP chip. At the 5M SNP density, ssGBLUP improved 0.95% in prediction accuracy compared to the 777K SNP chip. BayesR3 demonstrated an average increase of 2.08% in prediction accuracy at the 140K SNP chip compared to the 50K SNP chip. However, the prediction accuracy of BayesR3 decreased by 1.02% at the 777K SNP chip in contrast to the 140K SNP chip.

**Table 4:**
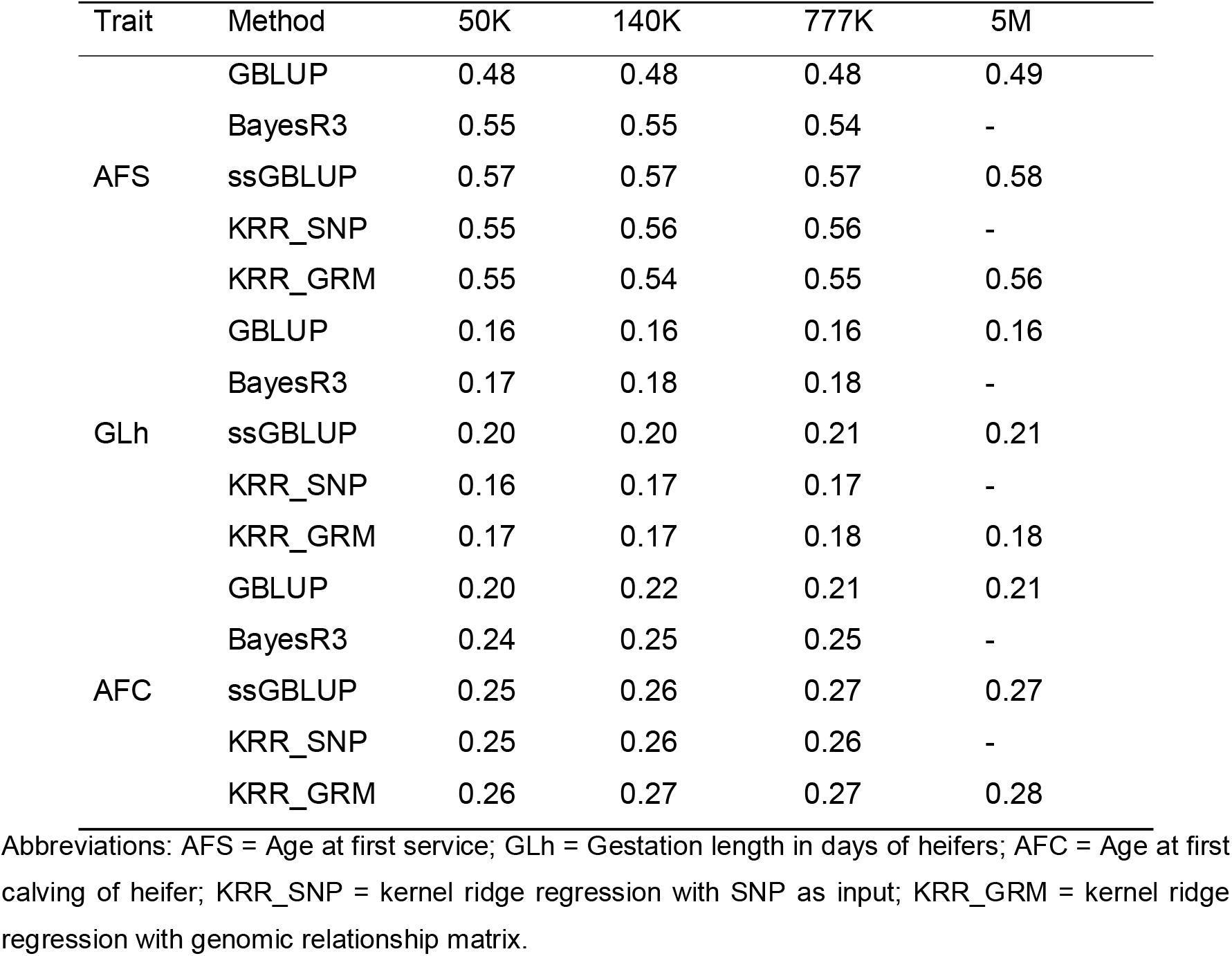
Prediction accuracy of GBLUP, ssGBLUP, BayesR3 and ML methods (KRR_SNP and KRR_GRM) of fertility traits.

Moreover, the efficiency of conventional methods was also influenced by the density of SNPs. For GBLUP, the running time increased by an average of 204.6% at 140K SNP chip compared to the 50K SNP chip. At the 777K SNP chip, the running time increased by an average of 251.5% compared to the 140K SNP chip. For ssGBLUP, the running time increased by an average of 214.4% at a SNP chip of 140K compared to the 50K SNP chip. At the 777K SNP chip, the running time increased by an average of 263.7% compared to the 140K SNP chip. Furthermore, at the 5M SNP chip, the running time increased by an average of 463.5% compared to the 777K SNP chip. As for BayesR3, the running time increased by an average of 160.1% at the 140K SNP chip compared to the 50K SNP chip. At the 777K SNP chip, the running time increased by an average of 367.2% compared to the 140K SNP chip. This indicates that as SNP density increases, the running time for conventional methods also significantly increases.

### Comparisons among ML Methods

In terms of prediction accuracy, the ML methods showed varying comparison results depending on the different types of input genotypes (Fig. 1). Firstly, when genotype data was inputted into the ML algorithms as an SNP matrix, KRR achieved the highest accuracy on the AFS, GLh, with average improvements of 0.82% and 17.02% compared to SVR, the second-best method, respectively. However, for the AFC trait, KRR’s accuracy was the second highest, lower than SVR. Across all three traits, RFR consistently exhibited the lowest accuracy. Secondly, when genotype data was inputted into the ML algorithms as an GRM, KRR achieves the highest accuracy on the GLh and AFC, with average improvements of 37.53% and 2.29% compared to the second-best method, respectively. However, for the AFS, KRR’s accuracy was the second highest. Lastly, when the genotypes are inputted in PCs form, the accuracy of the ML methods is significantly lower on the AFC and GLh compared to the other two formats. On the trait AFS, the accuracy of RFR was comparable to the other input formats, while other ML methods performed poorly.

The running time of three ML methods in all various genotype input forms displayed the same trend: KRR < SVR < RFR (Fig. 2). Specifically, KRR runs approximately 23-45 times faster (average of 35 times) than SVR and 130-361 times faster (average of 278 times) than RFR. It should be noted that we did not provide the test results for ML methods on the 5M SNP density in SNP form due to the large number of markers, which would be impractical in terms of both running time and memory consumption. Based on the results, we selected KRR_SNP and KRR_GRM for comparison with conventional methods.

### Comparisons of ML Methods with GBLUP, ssGBLUP and BayesR3

For prediction accuracy (Table 4), substantial differences were observed among the different methods. For AFS, ssGBLUP achieved the highest accuracy (0.57), which was 2.84%, 4.09%, 4.73%, and 18.65% higher than that of KRR_SNP, KRR_GRM, BayesR3, and GBLUP, respectively. For GLh, ssGBLUP also had the highest accuracy (0.23), outperforming BayesR3, KRR_GRM, KRR_SNP, and GBLUP by 16.04%, 17.14%, 23.0%, and 28.13%, respectively. For AFC, KRR_GRM yielded the highest accuracy (0.27), which was 2.86%, 5.19%, 9.46%, and 28.57% greater than that of ssGBLUP, KRR_SNP, BayesR3, and GBLUP, respectively. Furthermore, the unbiasedness of KRR_SNP and KRR_GRM was comparable to GBLUP, ssGBLUP, and BayesR3 on average (Table 5).

**Table 5:** Prediction unbiasedness of GBLUP, BayesR3 and ML methods (KRR_SNP and KRR_GRM) of fertility traits.

| Trait | Method | 50K | 140K | 777K | 5M |
| --- | --- | --- | --- | --- | --- |
| AFS | GBLUP | 0.97 | 0.98 | 0.98 | 1.0 |
|  | BayesR3 | 1.01 | 1.01 | 0.97 | - |
|  | ssGBLUP | 1.04 | 1.03 | 1.01 | 1.04 |
|  | KRR_SNP | 1.20 | 1.21 | 1.21 | - |
|  | KRR_GRM | 1.20 | 1.21 | 1.21 | 1.21 |
| cGLh | GBLUP | 0.96 | 0.99 | 0.99 | 0.96 |
|  | BayesR3 | 1.13 | 1.08 | 1.06 | - |
|  | ssGBLUP | 1.04 | 1.03 | 1.05 | 1.01 |
|  | KRR_SNP | 0.76 | 0.77 | 0.84 | - |
|  | KRR_GRM | 0.82 | 0.83 | 0.85 | 0.87 |
| AFC | GBLUP | 0.90 | 0.94 | 0.97 | 0.95 |
|  | BayesR3 | 0.98 | 0.97 | 0.95 | - |
|  | ssGBLUP | 0.98 | 0.98 | 1.00 | 1.02 |
|  | KRR_SNP | 0.95 | 0.98 | 1.04 | - |
|  | KRR_GRM | 0.98 | 0.99 | 0.99 | 1.01 |

Abbreviations: AFS = Age at first service; GLh = Gestation length in days of heifers; AFC = Age at first calving of heifer; KRR\_SNP = kernel ridge regression with SNP as input; KRR\_GRM = kernel ridge regression with genomic relationship matrix.

## Discussion

Breeding cost is a critical factor in animal breeding programs. Historically, most genotyping data in dairy cattle breeding were generated using SNP arrays, such as the 50K chip, but this approach is associated with relatively high costs. One of the main advantages for low-coverage sequencing is its low cost, which is approximately two-thirds lower than that of a standard 50K SNP chip. Although the low-coverage sequencing allows for the acquisition of whole-genome variants at a significantly lower cost, research on the application of low-coverage sequencing data in dairy cattle remains relatively limited. The present study evaluated the implementation of low-coverage sequencing data as a cost-efficient approach in Chinese Holstein heifers. Our results highlight the effectiveness of lcGWS data in breeding applications, and showed low-coverage sequencing data can achieve comparable or even higher accuracy than SNP arrays at a reduced cost. There have been extensive studies into the impact of SNP density on GP using GBLUP model, and results are different (Ober et al., 2012; van Binsbergen et al., 2015; Song et al., 2019; Tong et al., 2023; Xie et al., 2023). In this study, the effect of SNP density on GP accuracy was investigated using three fertility traits. The results indicate that the prediction accuracy of ssGBLUP remained largely consistent or slightly increased with SNP density, whereas the accuracy of other conventional methods (GBLUP, BayesR3) and the ML methods exhibited more complex fluctuations across different marker densities. Theoretically, the higher the density of SNPs, the greater likelihood of capturing causal mutations for phenotypic traits. Therefore, the prediction is much less constrained by the LD between the SNP and the causal mutation (Meuwissen and Goddard, 2010). In addition, as SNP density increases, it may capture more genetic variance, further increase accuracy. The imputation accuracy of low-coverage sequencing data is higher than that of SNP chip data (Zhang et al., 2023). Consequently, breeding workers can flexibly choose to either extract imputed SNP chip data or use the full whole-genome sequencing data for GS, depending on their specific breeding objectives. This flexibility represents another key advantage of low-coverage sequencing data.

In this study, ML methods shown highly GP accuracy when only using genotype data. A study highlighted that the ML methods make no assumptions about the SNP effect distribution and can flexibly capture the non-linear relationships, and they remain more efficient in the presence of larger datasets (Piles et al., 2021). Furthermore, the degree of dimensionality reduction from SNP to G matrix is higher, which may decrease SNP noise and improve prediction accuracy. It is also noteworthy that hyperparameter tuning has been performed in ML methods, and after multiple combinations of parameters, the accuracy can be improved. The KRR employed a cosine kernel, rather than a strictly linear kernel. This kernel normalizes genotype data, emphasizing angular similarity between samples rather than linear combinations. This normalization may enable KRR_GRM to better capture the complex genetic architecture of fertility traits. When using both genotype and pedigree data, our results showed that ssGBLUP achieved the highest prediction accuracy for AFS and GLh. This finding is largely consistent with previous studies (Teng et al., 2022; Wang et al., 2022), suggesting that including more genetic information has the potential to improve the accuracy of GP.

While this study demonstrates the practical value of lcGWS and ML, we acknowledge several limitations that offer avenues for future research. Firstly, while low-coverage sequencing offers key advantages in cost and flexibility, the current pipeline from raw sequencing data to imputed genotypes is multi-stepped and computationally complex, and would benefit from streamlining to facilitate broader adoption in routine breeding programs. Furthermore, the genomic dataset, while sufficient to validate the primary methods, is modest in size (n=3,683). This is particularly relevant for the ML models, as their capacity to capture complex non-linear patterns is often dependent on larger training populations. Consequently, the full potential of these ML methods compared to conventional models may be even greater with expanded datasets, such as by obtaining genotypes for all individuals with phenotype records. Secondly, our investigation was confined to specific ML algorithms. The rapid advancements in deep learning present a promising frontier for potentially improving genomic prediction further, especially for traits with complex genetic architectures. Lastly, our findings are based on fertility traits in a single breed. Future studies should therefore aim to validate these cost-effective strategies across other economically important traits and in different cattle breeds to confirm their broad applicability.

## Conclusion

In summary, this study demonstrates the effectiveness of lcGWS data in cattle breeding applications, and low-coverage sequencing data can achieve comparable or even higher accuracy than SNP arrays at a reduced cost. Our results also showed the ML’s potential to enhance GP accuracy for fertility traits. It also serves as a reference to improve breeding efficiency in livestock.

## Ethics approval

Not applicable.

## Data and model availability statement

The datasets used in the current study are available from the corresponding author on reasonable request.

## Declaration of generative AI and AI-assisted technologies in the writing process

The authors did not use any artificial intelligence-assisted technologies in the writing process.

## Acknowledgements

We thank the high-performance computing platform of Northwest A&F University and Xian advanced computing center for providing computing resources.

## References

Alves, K., Brito, L.F., Schenkel, F.S., 2023. Genomic prediction of fertility and calving traits in Holstein cattle based on models including epistatic genetic effects. J Anim Breed Genet 140, 568–581. doi:10.1111/jbg.12810.

An, B., Liang, M., Chang, T., Duan, X., Du, L., Xu, L., Zhang, L., Gao, X., Li, J., Gao, H., 2021. KCRR: a nonlinear machine learning with a modified genomic similarity matrix improved the genomic prediction efficiency. Brief Bioinform 22, bbab132. doi:10.1093/bib/bbab132.

Aponte, P.F.C., Carneiro, P.L.S., Araujo, A.C., Pedrosa, V.B., Fotso-Kenmogne, P.R., Silva, D.A., Miglior, F., Schenkel, F.S., Brito, L.F., 2024. Investigating the genomic background of calving-related traits in Canadian Jersey cattle. J Dairy Sci 107, 11195–11213. doi:10.3168/jds.2024-24768.

Breen, E.J., MacLeod, I.M., Ho, P.N., Haile-Mariam, M., Pryce, J.E., Thomas, C.D., Daetwyler, H.D., Goddard, M.E., 2022. BayesR3 enables fast MCMC blocked processing for largescale multi-trait genomic prediction and QTN mapping analysis. Commun Biol 5, 661. doi:10.1038/s42003-022-03624-1.

Breiman, L., 2001. Random forests. Machine Learning 45, 5–32. doi:Doi 10.1023/A:1010933404324.

Chafai, N., Hayah, I., Houaga, I., Badaoui, B., 2023. A review of machine learning models applied to genomic prediction in animal breeding. Front Genet 14, 1150596. doi:10.3389/fgene.2023.1150596.

Chang, C.C., Chow, C.C., Tellier, L.C., Vattikuti, S., Purcell, S.M., Lee, J.J., 2015. Second-generation PLINK: rising to the challenge of larger and richer datasets. Gigascience 4, 7. doi:10.1186/s13742-015-0047-8.

Chen, S., Zhou, Y., Chen, Y., Gu, J., 2018. fastp: an ultra-fast all-in-one FASTQ preprocessor. Bioinformatics 34, i884–i890. doi:10.1093/bioinformatics/bty560.

Drucker H B.C.J., Kaufman L, Alex S, Vapink V, 1997. Support vector regression machines. Advances In Neural Information Processing Systems 28, 779–784.

Ehret, A., Hochstuhl, D., Gianola, D., Thaller, G., 2015. Application of neural networks with back-propagation to genome-enabled prediction of complex traits in Holstein-Friesian and German Fleckvieh cattle. Genet Sel Evol 47, 22. doi:10.1186/s12711-015-0097-5.

Erbe, M., Hayes, B.J., Matukumalli, L.K., Goswami, S., Bowman, P.J., Reich, C.M., Mason, B.A., Goddard, M.E., 2012. Improving accuracy of genomic predictions within and between dairy cattle breeds with imputed high-density single nucleotide polymorphism panels. J Dairy Sci 95, 4114–4129. doi:10.3168/jds.2011-5019.

Garcia-Ruiz, A., Cole, J.B., VanRaden, P.M., Wiggans, G.R., Ruiz-Lopez, F.J., Van Tassell, C.P., 2016. Changes in genetic selection differentials and generation intervals in US Holstein dairy cattle as a result of genomic selection. Proc Natl Acad Sci U S A 113, E3995–4004. doi:10.1073/pnas.1519061113.

Habier, D., Fernando, R.L., Kizilkaya, K., Garrick, D.J., 2011. Extension of the bayesian alphabet for genomic selection. BMC Bioinformatics 12, 186. doi:10.1186/1471-2105-12-186.

Hong, J.K., Kim, Y.M., Cho, E.S., Lee, J.B., Kim, Y.S., Park, H.B., 2024. Application of deep learning with bivariate models for genomic prediction of sow lifetime productivity-related traits. Anim Biosci 37, 622–630. doi:10.5713/ab.23.0264.

Lee, H.J., Lee, J.H., Gondro, C., Koh, Y.J., Lee, S.H., 2023. deepGBLUP: joint deep learning networks and GBLUP framework for accurate genomic prediction of complex traits in Korean native cattle. Genet Sel Evol 55, 56. doi:10.1186/s12711-023-00825-y.

Li, H., Durbin, R., 2010. Fast and accurate long-read alignment with Burrows-Wheeler transform. Bioinformatics 26, 589–595. doi:10.1093/bioinformatics/btp698.

Li, H., Handsaker, B., Wysoker, A., Fennell, T., Ruan, J., Homer, N., Marth, G., Abecasis, G., Durbin, R., Genome Project Data Processing, S., 2009. The Sequence Alignment/Map format and SAMtools. Bioinformatics 25, 2078–2079. doi:10.1093/bioinformatics/btp352.

Liang, M., Cao, S., Deng, T., Du, L., Li, K., An, B., Du, Y., Xu, L., Zhang, L., Gao, X., Li, J., Guo, P., Gao, H., 2023. MAK: a machine learning framework improved genomic prediction via multi-target ensemble regressor chains and automatic selection of assistant traits. Brief Bioinform 24, bbad043. doi:10.1093/bib/bbad043.

Liang, M., Chang, T., An, B., Duan, X., Du, L., Wang, X., Miao, J., Xu, L., Gao, X., Zhang, L., Li, J., Gao, H., 2021. A Stacking Ensemble Learning Framework for Genomic Prediction. Front Genet 12, 600040. doi:10.3389/fgene.2021.600040.

Liu, H., Xing, K., Jiang, Y., Liu, Y., Wang, C., Ding, X., 2022. Using Machine Learning to Identify Biomarkers Affecting Fat Deposition in Pigs by Integrating Multisource Transcriptome Information. J Agric Food Chem 70, 10359–10370. doi:10.1021/acs.jafc.2c03339.

Martin, A.R., Atkinson, E.G., Chapman, S.B., Stevenson, A., Stroud, R.E., Abebe, T., Akena, D., Alemayehu, M., Ashaba, F.K., Atwoli, L., Bowers, T., Chibnik, L.B., Daly, M.J., DeSmet, T., Dodge, S., Fekadu, A., Ferriera, S., Gelaye, B., Gichuru, S., Injera, W.E., James, R., Kariuki, S.M., Kigen, G., Koenen, K.C., Kwobah, E., Kyebuzibwa, J., Majara, L., Musinguzi, H., Mwema, R.M., Neale, B.M., Newman, C.P., Newton, C., Pickrell, J.K., Ramesar, R., Shiferaw, W., Stein, D.J., Teferra, S., van der Merwe, C., Zingela, Z., Neuro, G.A.P.P.S.T., 2021. Low-coverage sequencing cost-effectively detects known and novel variation in underrepresented populations. Am J Hum Genet 108, 656–668. doi:10.1016/j.ajhg.2021.03.012.

Meuwissen, T., Goddard, M., 2010. Accurate prediction of genetic values for complex traits by whole-genome resequencing. Genetics 185, 623–631. doi:10.1534/genetics.110.116590.

Meuwissen, T.H., Hayes, B.J., Goddard, M.E., 2001. Prediction of total genetic value using genome-wide dense marker maps. Genetics 157, 1819–1829. doi:10.1093/genetics/157.4.1819.

Misztal, I., S. Tsuruta, D. A. L. Lourenco, Y. Masuda, I. Aguilar, A. Legarra, Vitezica, Z., 2018. Manual for BLUPF90 family programs. http://nce.ads.uga.edu/wiki/doku.php?id=documentation

Ober, U., Ayroles, J.F., Stone, E.A., Richards, S., Zhu, D., Gibbs, R.A., Stricker, C., Gianola, D., Schlather, M., Mackay, T.F., Simianer, H., 2012. Using whole-genome sequence data to predict quantitative trait phenotypes in Drosophila melanogaster. PLoS Genet 8, e1002685. doi:10.1371/journal.pgen.1002685.

Pedregosa, F., Varoquaux, G., Gramfort, A., Michel, V., Thirion, B., Grisel, O., Blondel, M., Prettenhofer, P., Weiss, R., Dubourg, V.J.t.J.o.m.L.r., 2011. Scikit-learn: Machine learning in Python. Journal ofMachine Learning Research 12, 2825–2830.

Piles, M., Bergsma, R., Gianola, D., Gilbert, H., Tusell, L., 2021. Feature Selection Stability and Accuracy of Prediction Models for Genomic Prediction of Residual Feed Intake in Pigs Using Machine Learning. Front Genet 12, 611506. doi:10.3389/fgene.2021.611506.

Ronneburg, T., Zan, Y., Honaker, C.F., Siegel, P.B., Carlborg, O., 2023. Low-coverage sequencing in a deep intercross of the Virginia body weight lines provides insight to the polygenic genetic architecture of growth: novel loci revealed by increased power and improved genome-coverage. Poult Sci 102, 102203. doi:10.1016/j.psj.2022.102203.

Schaeffer, L.R., 2006. Strategy for applying genome-wide selection in dairy cattle. J Anim Breed Genet 123, 218–223. doi:10.1111/j.1439-0388.2006.00595.x.

Song, H., Ye, S., Jiang, Y., Zhang, Z., Zhang, Q., Ding, X., 2019. Using imputation-based whole-genome sequencing data to improve the accuracy of genomic prediction for combined populations in pigs. Genet Sel Evol 51, 58. doi:10.1186/s12711-019-0500-8.

Teng, J.-y., Ye, S.-p., Gao, N., Chen, Z.-t., Diao, S.-q., Li, X.-j., Yuan, X.-l., Zhang, H., Li, J.-q., Zhang, X.-q., Zhang, Z., 2022. Incorporating genomic annotation into single-step genomic prediction with imputed whole-genome sequence data. Journal of Integrative Agriculture 21, 1126–1136. doi:10.1016/s2095-3119(21)63813-3.

Tong, X., Chen, D., Hu, J., Lin, S., Ling, Z., Ai, H., Zhang, Z., Huang, L., 2023. Accurate haplotype construction and detection of selection signatures enabled by high quality pig genome sequences. Nat Commun 14, 5126. doi:10.1038/s41467-023-40434-3.

van Binsbergen, R., Calus, M.P., Bink, M.C., van Eeuwijk, F.A., Schrooten, C., Veerkamp, R.F., 2015. Genomic prediction using imputed whole-genome sequence data in Holstein Friesian cattle. Genet Sel Evol 47, 71. doi:10.1186/s12711-015-0149-x.

VanRaden, P.M., Van Tassell, C.P., Wiggans, G.R., Sonstegard, T.S., Schnabel, R.D., Taylor, J.F., Schenkel, F.S., 2009. Invited review: reliability of genomic predictions for North American Holstein bulls. J Dairy Sci 92, 16–24. doi:10.3168/jds.2008-1514.

Vovk, V., 2013. Kernel Ridge Regression. In Empirical Inference (ed. Schölkopf, B., Luo, Z. and Vovk, V.), Springer Berlin Heidelberg, Berlin, Heidelberg, p. 105–116. doi:10.1007/978-3-642-41136-6_11.

Wang, K., Abid, M.A., Rasheed, A., Crossa, J., Hearne, S., Li, H., 2023a. DNNGP, a deep neural network-based method for genomic prediction using multi-omics data in plants. Mol Plant 16, 279–293. doi:10.1016/j.molp.2022.11.004.

Wang, X., Shi, S., Ali Khan, M.Y., Zhang, Z., Zhang, Y., 2024. Improving the accuracy of genomic prediction in dairy cattle using the biologically annotated neural networks framework. J Anim Sci Biotechnol 15, 87. doi:10.1186/s40104-024-01044-1.

Wang, X., Shi, S., Wang, G., Luo, W., Wei, X., Qiu, A., Luo, F., Ding, X., 2022. Using machine learning to improve the accuracy of genomic prediction of reproduction traits in pigs. J Anim Sci Biotechnol 13, 60. doi:10.1186/s40104-022-00708-0.

Wang, Z., Ma, H., Li, H., Xu, L., Li, H., Zhu, B., Hay, E.H., Xu, L., Li, J., 2023b. Multi-trait genomic predictions using GBLUP and Bayesian mixture prior model in beef cattle. Animal Research and One Health 1, 17–29. doi:10.1002/aro2.13.

Xiang, T., Li, T., Li, J., Li, X., Wang, J., 2023. Using machine learning to realize genetic site screening and genomic prediction of productive traits in pigs. FASEB J 37, e22961. doi:10.1096/fj.202300245R.

Xie, L., Qin, J., Rao, L., Cui, D., Tang, X., Chen, L., Xiao, S., Zhang, Z., Huang, L., 2023. Genetic dissection and genomic prediction for pork cuts and carcass morphology traits in pig. J Anim Sci Biotechnol 14, 116. doi:10.1186/s40104-023-00914-4.

Yan, J., Wang, X., 2023. Machine learning bridges omics sciences and plant breeding. Trends Plant Sci 28, 199–210. doi:10.1016/j.tplants.2022.08.018.

Yang, B., Li, Y., Li, Q., Liu, S., 2024. High-throughput and cost-effective genotyping by low-coverage whole genome sequencing with genotype imputation in Pacific oyster, Crassostrea gigas. Aquaculture 591 doi:10.1016/j.aquaculture.2024.741134.

Yang, J., Lee, S.H., Goddard, M.E., Visscher, P.M., 2011. GCTA: a tool for genome-wide complex trait analysis. Am J Hum Genet 88, 76–82. doi:10.1016/j.ajhg.2010.11.011.

Yang, R., Guo, X., Zhu, D., Tan, C., Bian, C., Ren, J., Huang, Z., Zhao, Y., Cai, G., Liu, D., Wu, Z., Wang, Y., Li, N., Hu, X., 2021. Accelerated deciphering of the genetic architecture of agricultural economic traits in pigs using a low-coverage whole-genome sequencing strategy. Gigascience 10, giab048. doi:10.1093/gigascience/giab048.

Yin, L., Zhang, H., Zhou, X., Yuan, X., Zhao, S., Li, X., Liu, X., 2020. KAML: improving genomic prediction accuracy of complex traits using machine learning determined parameters. Genome Biol 21, 146. doi:10.1186/s13059-020-02052-w.

Zhang, H., Yin, L., Wang, M., Yuan, X., Liu, X., 2019. Factors Affecting the Accuracy of Genomic Selection for Agricultural Economic Traits in Maize, Cattle, and Pig Populations. Front Genet 10, 189. doi:10.3389/fgene.2019.00189.

Zhang, Z., Wang, A., Hu, H., Wang, L., Gong, M., Yang, Q., Liu, A., Li, R., Zhang, H., Zhang, Q., Shah, A.M., Wang, X., Wang, Y., Liu, Q., Gao, L., Zhang, Z., Wang, C., Ma, Y., Cai, Y., Jiang, Y., 2023. The efficient phasing and imputation pipeline of low - coverage whole genome sequencing data using a high - quality and publicly available reference panel in cattle. Animal Research and One Health 1, 4–16. doi:10.1002/aro2.8.

